# Proteomic signatures of podocyte injury are reflected in urinary extracellular vesicles in pediatric nephrotic syndrome

**DOI:** 10.1101/2025.08.19.670903

**Authors:** Tyler T. Cooper, Owen Hovey, Gilles A. Lajoie, Christopher R.J. Kennedy, Dylan Burger, Robert L. Myette

**Author notes:** **Correspondence:** Robert L. Myette, Kidney Research Center, Ottawa Hospital Research Institute, Department of Cellular and Molecular Medicine, University of Ottawa, 2518-451 Smyth Rd. Ottawa, ON, Canada.

## Abstract

Idiopathic nephrotic syndrome (NS) is a common glomerulopathy in children and presents with significant proteinuria. There are no reliable clinical or biochemical markers of disease relapse, or prognosis. Extracellular vesicles (EVs) are small, membrane-bound biological effectors released from stressed cells. We previously showed increases in podocyte-specific urinary EVs from children with disease relapse in NS, with numbers returning to near-zero in remission. Herein, we have expanded this work to evaluate puromycin aminonucleoside (PAN) injury by characterizing proteomic signatures of podocytes and their EVs *in vitro*. In addition, we performed data-independent proteomic analysis (DIA) to characterize changes in signatures of EVs from pediatric patients with active NS versus remission. Our key findings reveal PAN-injured podocytes increase large EV (LEV) secretion *in vitro*; moreover, DIA uncovered changes in cellular and LEV proteomes that were also observed in urinary LEVs from patients with active disease. Urinary LEV proteomes from children with active NS were significantly different than those in remission, highlighted by 645 and 240 unique proteins associated with disease or remission, respectively. This foundational work provides the impetus for a larger, prospective biomarker study aimed at identifying EV-specific proteins associated with relapse versus remission.

## Introduction

Idiopathic nephrotic syndrome (NS) is a common glomerulopathy in children affecting up to 1:5000 children and presents with significant proteinuria, hypoalbuminemia, and edema [1]. Podocytopathy underlies the etiology of NS and can be confirmed by observing podocyte effacement following kidney biopsy and pathological examination [2]. Current global standards for elucidating the pathophysiology of NS in humans requires the use of both *in vivo* and *in vitro* models that mimic podocyte injury. Chemical insult using puromycin aminonucleoside (PAN) is a widely accepted approach to model podocytopathy in immortalized human podocytes *in vitro* and is a model of nephrosis in rat models *in vivo* [3–5]. PAN is thought to act via an oxidative stress mechanism which in turn ‘mimics’ renal features of NS by eliciting podocyte injury [6].

Extracellular vesicles (EVs) are membrane-encapsulated vectors released from almost all cell types during times of cellular stress to transfer a diverse range of bioactive cargo [7]. These vesicles contain DNA, miRNA, metabolites, and protein which allow for complex cell-to-cell signaling under both physiological and pathological conditions [7]. A population of large EVs (LEVs), historically referred to as microparticles, are ∼100 to 1000 nm in diameter, and are derived predominantly from budding of the plasma membrane [8], retaining characteristics of the parent cell of origin. Recently, we showed that podocyte-specific urinary LEVs from children with NS were increased during times of relapse and returned to near-zero levels during remission [9]. However, comprehensive proteomic characterization of urinary LEVs in pediatric NS is lacking, and their potential as biomarkers for diagnosis and therapeutic response is currently unrealized.

Proteomics is the large-scale study of protein functions, compositions, and interactions, offering unparalleled insights into cellular processes and mechanisms of disease. This approach is particularly valuable for studying EVs, given their diverse protein cargo and multifaceted roles in intercellular communication [10]. Advances in mass spectrometry-based proteomics, including data-independent acquisition (DIA) methodologies, now enable high-resolution, quantitative analysis of EV proteomes isolated from cell culture or biofluids. These techniques enable unbiased investigations that reveal molecular signatures harbored by tissue-specific EVs, with significant implications for understanding health and disease [11].

Given that NS is a disease of the podocyte and that podocyte-specific LEVs are increased in the urine of children with NS during times of relapse [9], we sought to better characterize LEV levels *in vitro* using conditionally immortalized human podocytes (hPods). We observed that the production of LEVs in hPods increased following treatment with PAN. To understand the molecular impact of PAN treatment on podocyte LEVs, we performed proteomic analyses of hPod-derived LEVs and the hPods themselves *in vitro* following PAN exposure. We then compared this *in vitro* model with urinary LEVs isolated from children with NS during both disease and remission, highlighting shared features and significant differences between the cellular model and clinical samples. Herein, we provide the first comprehensive proteomic characterization of urinary LEVs from children with NS, uncovering a diverse array of proteins from both glomerular and tubular origin. Our work provides foundational knowledge on the molecular composition of LEVs in pediatric NS, informs future efforts in biomarker discovery, and offers potential for future therapeutic monitoring.

## Methodology

### Podocyte Culture

hPods were obtained with permission of Prof. Moin Saleem (University of Bristol, Bristol, UK) [3]. Cells were grown in RPMI-1640 medium supplemented with 10% Fetal Bovine Serum (Invitrogen, Carlsbad, CA) and penicillin-streptomycin (1:100; Invitrogen). Podocytes were proliferated at 33°C in the presence of 10 U/mL Recombinant Human Interferon Gamma (γ-IFN; Invitrogen). For induction of podocyte differentiation, cells were maintained at 38°C for 13-14 days in the absence of γ-IFN, with the experiment performed, in all cases, between days 13-14. PAN (25 μg/mL; Cayman Chemicals, Ann Arbour, MI) is a well-characterized podocyte toxin that models NS in the laboratory. PAN dose was selected based on dose-response experiments and occurred for 24-hours, as indicated. After treatment, the media was collected, and LEVs were isolated as described. Cell plates were washed with ice cold PBS immediately after media collection and were frozen at −80°C until protein isolation. Protein concentration was determined using the DC protein assay (Bio-Rad, Hercules, CA, USA, Catalog 5000111).

### Pediatric Nephrotic Syndrome Urine

Our prospective study enrolling children with NS is ongoing at the Children’s Hospital of Eastern Ontario (REB #18/168X). Urine was obtained from 8 children with NS during both the relapse and remission state. Remission was defined as a urine protein to creatinine ratio of < 0.05 g/mmol. Enrolled children have NS, as defined clinically, and when performed, confirmed by biopsy. Not all children in this study had a biopsy. Those that did not were presumed to have NS associated with minimal change lesions (**Supplemental Table 1**). Urine samples were freshly obtained and were centrifuged at 2,500 x g for 10 minutes within 4 hours of voiding. Thereafter, urine was frozen at −80 C until used. There were no freeze/thaw events. There were 4 relapse samples, and 4 remission samples evaluated in this cross-sectional study. We analyzed samples from children with steroid sensitive and steroid resistant disease. If dependent on steroids, they were also deemed steroid sensitive. Those who had no response to steroids, or who had steroids discontinued and were escalated to a second-line agent, were deemed steroid resistant (**Supplemental Table 1**).

### Extracellular Vesicles Isolation

LEVs were isolated using sequential centrifugation as described previously [9, 12, 13]. Briefly, cell culture media was centrifuged at 2,500 x *g* for 10 minutes, the supernatant was then collected and centrifuged at 20,000 x *g* for 20 minutes to pellet LEVs. These LEVs were resuspended in sterile, pre-filtered (0.1 μm) phosphate buffered saline (PBS) and frozen at −80°C until analysis. Urinary LEVs were isolated similarly. Briefly, urine was thawed and vortexed. Subsequently, the urine was aliquoted and centrifuged at 20,000 x g for 20 minutes to pellet LEVs. Urinary LEVs were resuspended in pre-filtered PBS, and frozen prior to proteomic analysis.

### Nanoparticle Tracking Analysis

NTA was performed as previously described [9]. Briefly, the ZetaView PMX110 Multiple Parameter Particle Tracking Analyzer (Particle Metrix, Meerbusch, Germany) was used. We used size-mode analysis with ZetaView software (version 8.02.28) following calibration with polystyrene beads (105 and 500 nm). Samples were analyzed at 11 camera positions with 2-second video length at 21°C. LEVs were quantified per experiment and expressed as relative to control.

### Liquid Chromatography and Tandem Mass Spectrometry

The MS/MS spectra were obtained using the protocol previously reported by our group [11]. Briefly, 10 μg of EV or 50 μg of cell lysates in radioimmunoprecipitation assay buffer (150 mM NaCl, 1% Triton X-100, 0.5% sodium deoxycholate, 0.1% SDS, 50 mM Tris, pH 8.0) were isolated by single-pot protein precipitation (SP3) using 50% Ethanol and proteolysis (50 mM Ammonium Bicarbonate, 1:50 TrypLysC, 18 hours, 37°C) and analyzed on a Thermo Eclipse using Gas-phase Fractionation operating in Data-Independent Acquisition (GPF-DIA). Spectral libraries were first searched in DIA-NN against the human proteome before quantitative analysis using label-free quantification (MaxLFQ). The software parameters were set to a precursor mass tolerance of 20 ppm and a fragment ion mass tolerance of 10 ppm. The search parameters allowed for up to 2 missed cleavages and a maximum of 2 post-translational modifications per peptide. Missing value imputation, transformations, and proteomic data visualizations were performed using in-house Python code. Differentially expressed proteins were defined as follows: abs|log2 Fold Change| > 0.6 & log10(p-value) > 1.30.

### Data Analysis

Continuous variables were summarized as mean (±standard deviation, SD) if normally distributed. When non-normally distributed, medians with interquartile ranges were reported. P-values were generated using student t-tests or one-way analysis of variance with multiple comparisons which were corrected using Tukey’s method when normally distributed, and Kruskal Wallis and U-test when non-normally distributed. Tests were performed using Prism (version 9.4.1) and Python (version 3.13.1).

## Results

### PAN-induced injury drives distinct proteomic changes in immortalized human podocytes

This study evaluated the impact of PAN (25 μg/mL) on changes to the hPod proteome (**Figure 1A**). Overall, we identified and quantified >5000 proteins with unique proteomic profiles for each experimental condition. Importantly, proteomic changes did not coincide with an increase or decrease in cell number, as measured using total cellular protein levels between each condition (**Figure 1B**). Principal component analysis (PCA) revealed distinct proteomic profiles between both groups (**Figure 1C**). We then sought to examine differentially expressed proteins (DEPs, defined as: abs|log2 Fold Change| > 0.6 & log10(p-value) > 1.30) unique to PAN-injury. We observed 56 DEPs were upregulated in PAN-induced hPod injury relative to control, with 111 downregulated (**Figure 1D**).

**Figure 1.**
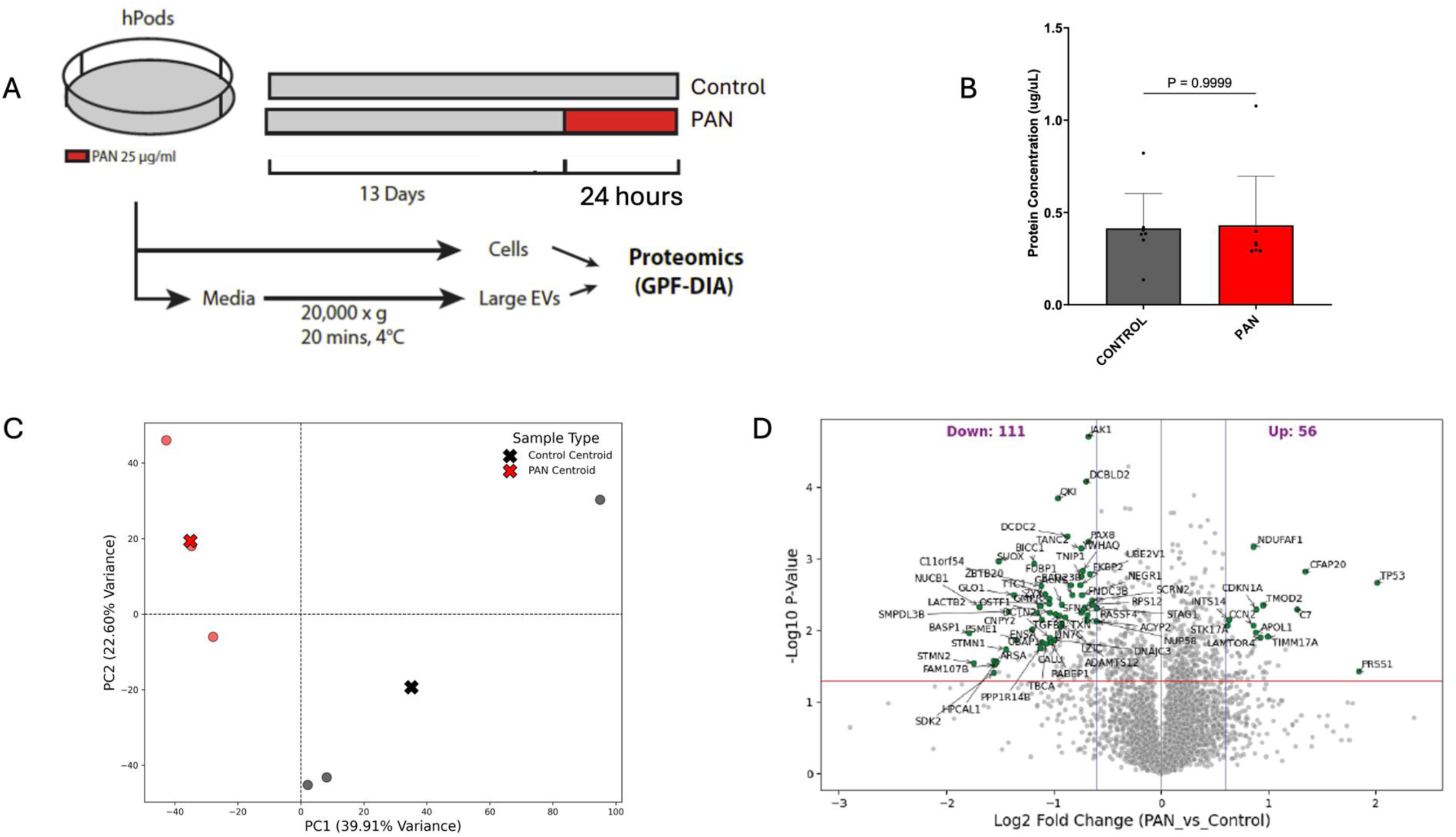
A. Cell culture (human podocytes, hPods) conditions and schematic of proteomic workflow. PAN, puromycin aminonucleoside; GPF-DIA, gas-phase fractionation with data-independent acquisition. B. Protein concentration from cell culture experiments, compared across both treatment conditions. C. Principal component analysis for cell culture analysis. D. Volcano plot comparing DEPs between PAN and control. This revealed an upregulation in 56 DEPs after PAN treatment, while 111 DEPs were downregulated.

### Proteomic transformations following PAN-injury are related to EV production and stress

To understand how DEPs confer phenotypic changes, we next used Gene Set Enrichment Analysis (GSEA) to uncover pathways differentially enriched between PAN-treated and control hPods. Initially, we focused on cellular compartments within PAN-injured hPods revealing a significant enrichment in proteins related to DNA stress responses and production of EVs (**Supplemental Figure 1A, B**).

### PAN treatment leads to an increase in hPod LEV formation and release and alters the LEV proteome

Following 24-hours of PAN treatment, hPod LEV release increased ∼3-fold (control 1.1, PAN 2.7, mean difference 1.6, 95% confidence interval 0.4 to 2.8; P=0.0045; **Figure 2A**). There was no apparent impact on LEV size under each treatment condition (**Figure 2B**).

**Figure 2.**
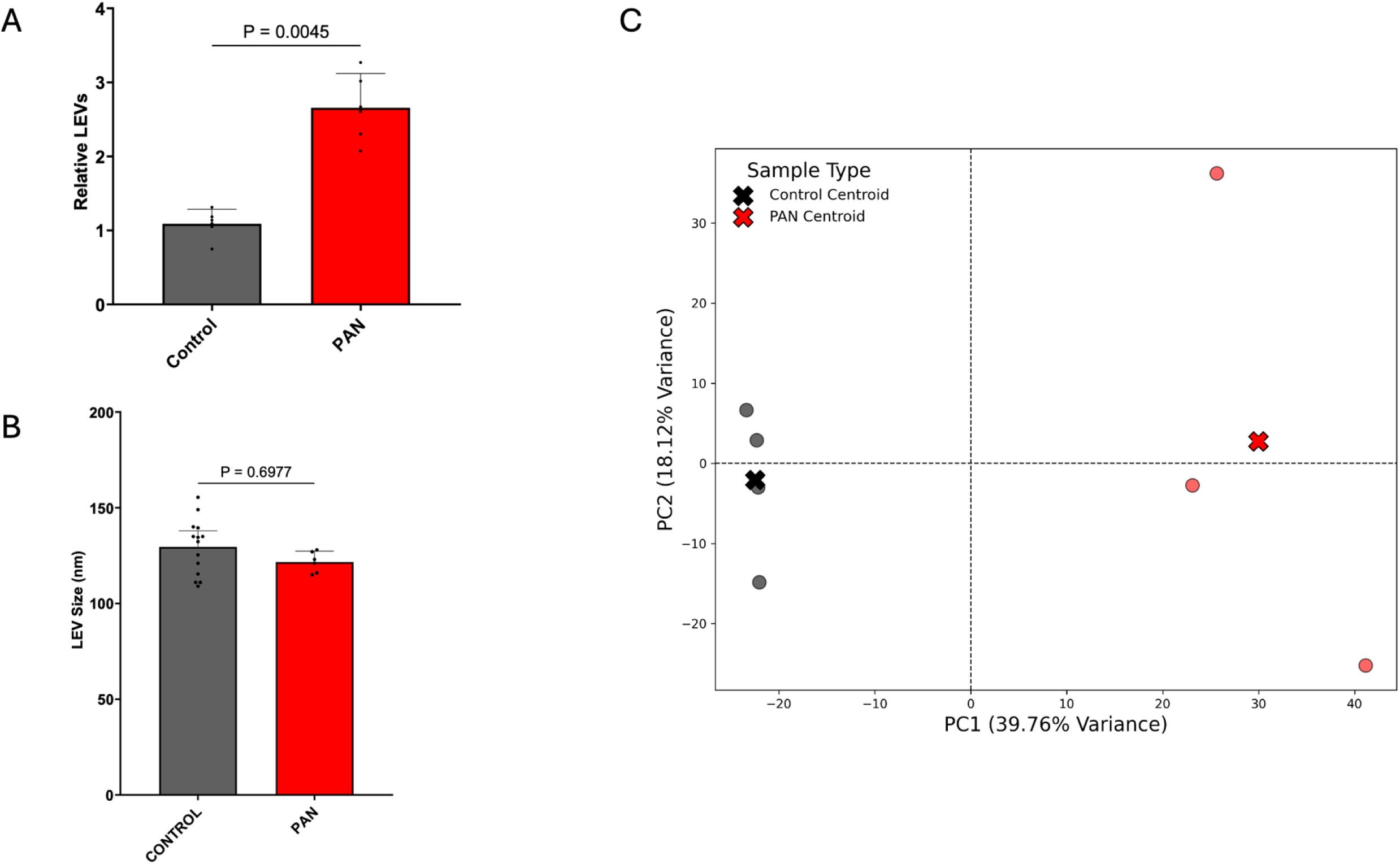
A. Relative large extracellular vesicle (LEV) production across all experimental conditions. Puromycin aminonucleoside (PAN) treatment led to a significant increase in LEV production (P=0.0045). B. There were no significant differences in the size of LEVs produced under both experimental conditions. C. Principal component analysis for LEV DEPs across each experimental condition.

Specifically, control hPod LEVs had a mean size of 129 nm compared to the mean size of PAN treated hPod LEVs, which was 122 nm (mean difference 7 nm, 95% confidence interval −8.4 to 24.2, P>0.05). Similar to cellular proteomes, LEVs have differing proteomes between experimental conditions, as evidenced by PCA analysis (**Figure 2C**). Importantly, EV markers (Albumin, Tissue Factor, CD9, and Programmed Cell Death 6 Interacting Protein (PDCD6IP or ALIX)) were not different between each experimental condition. This confirms consistent LEV isolation and supports our hypothesis that proteomic cargo in LEVs is unique to experimental conditions (**Supplemental Figure 2A**).

Next, we sought to examine LEV DEPs present between PAN-treated and control hPods. PAN-induced injury led to upregulation of 54 unique proteins compared to 138 downregulated proteins under control conditions (**Figure 3**). In support of these observations, GSEA analysis using Reactome and Jensen Compartments annotation revealed a strong enrichment in PAN LEVs of proteins related to Translation, Elongation and Endoplasmic Reticulum, respectively (**Supplemental Figure 3A-B**).

**Figure 3.**
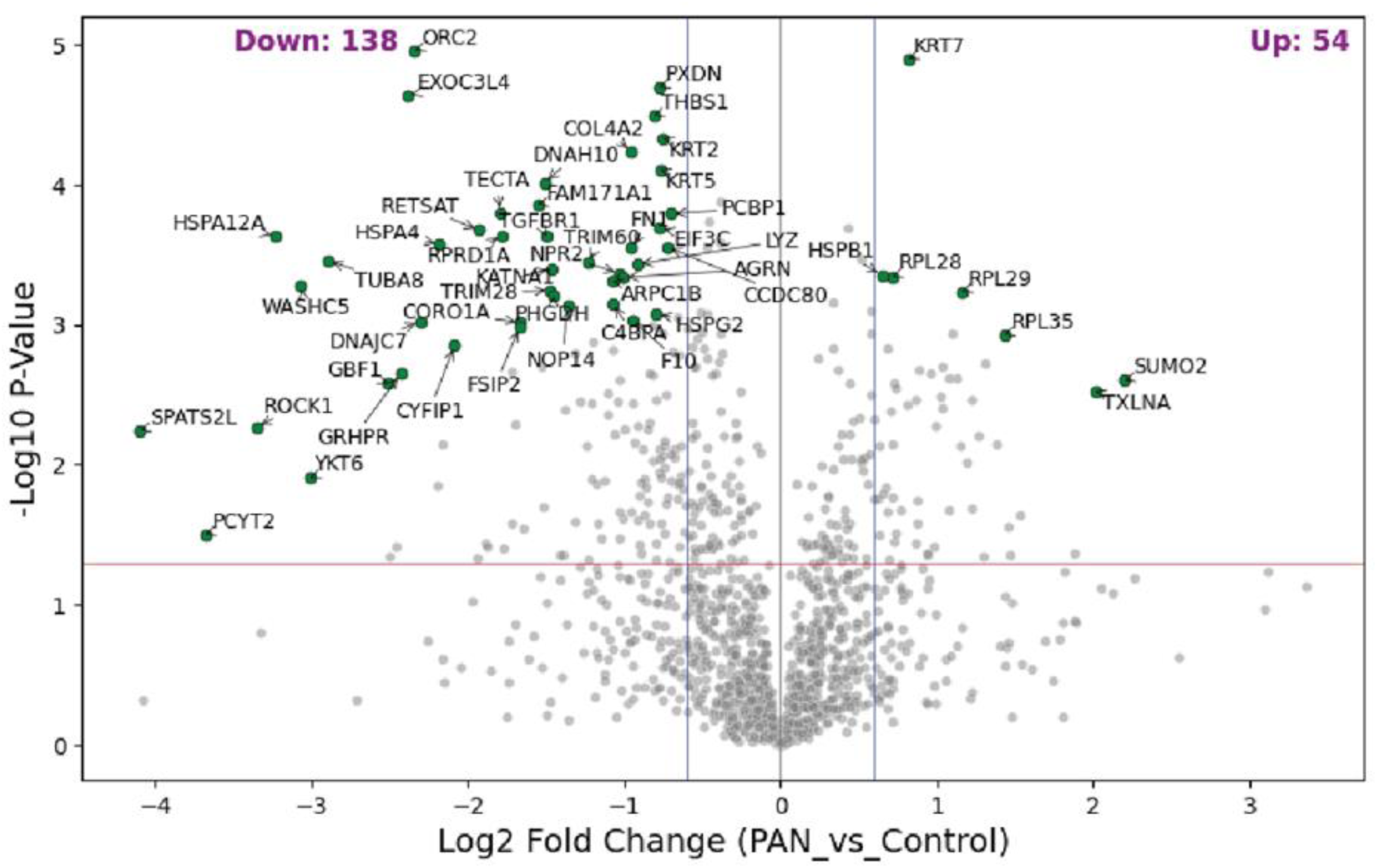
A. Volcano plot comparing differentially expressed proteins (DEPs) from large extracellular vesicles (LEVs) between puromycin aminonucleoside (PAN) and control. There were 54 upregulated DEPs in LEVs from human podocytes (hPods) following treatment with PAN, compared to 138 downregulated, relative to control.

### LEVs from children with NS harbor distinct proteomes reflective of disease status

Building on our *in vitro* analysis of the hPod cellular and LEV proteomes, and our recent work revealing an increase in urinary podocyte-specific LEVs in children with NS during relapse [9], we investigated whether human urinary LEVs from children with NS exhibit significant proteomic differences between disease and remission states. Patient characteristics are summarized in **Supplemental Table 1**. Our analysis identified approximately ∼2600 LEV proteins across both conditions, including 645 uniquely associated with disease and 240 proteins unique to remission (**Figure 4A**). Jensen Compartment Analysis revealed that these proteins were predominantly localized to extracellular organelles, extracellular vesicles and exosomes (**Figure 4A**). Proteins unique to disease were significantly associated with an immunological response, vascular endothelial growth factor (VEGF) signaling, and vesicle-mediated transport, whereas proteins unique to remission LEVs were associated with cell-cell adhesion and extracellular matrix reorganization (**Supplemental Figure 4A-B**). PCA revealed distinct proteomic profiles between disease and remission, highlighting the different LEV proteomes associated with each condition. (**Figure 4B**). To confirm reproducible LEV isolation from urine, we assessed common EV markers (CD9/81, PDCD6IP, and TSG101) and found no significant difference between conditions (**Figure 4C**). However, higher levels of albumin (P<0.05) and tissue factor (P<0.05) in disease-state LEVs confirmed their association with glomerulopathy, and for albumin, likely reflects non-specific binding of albumin in the context of proteinuria.

**Figure 4.**
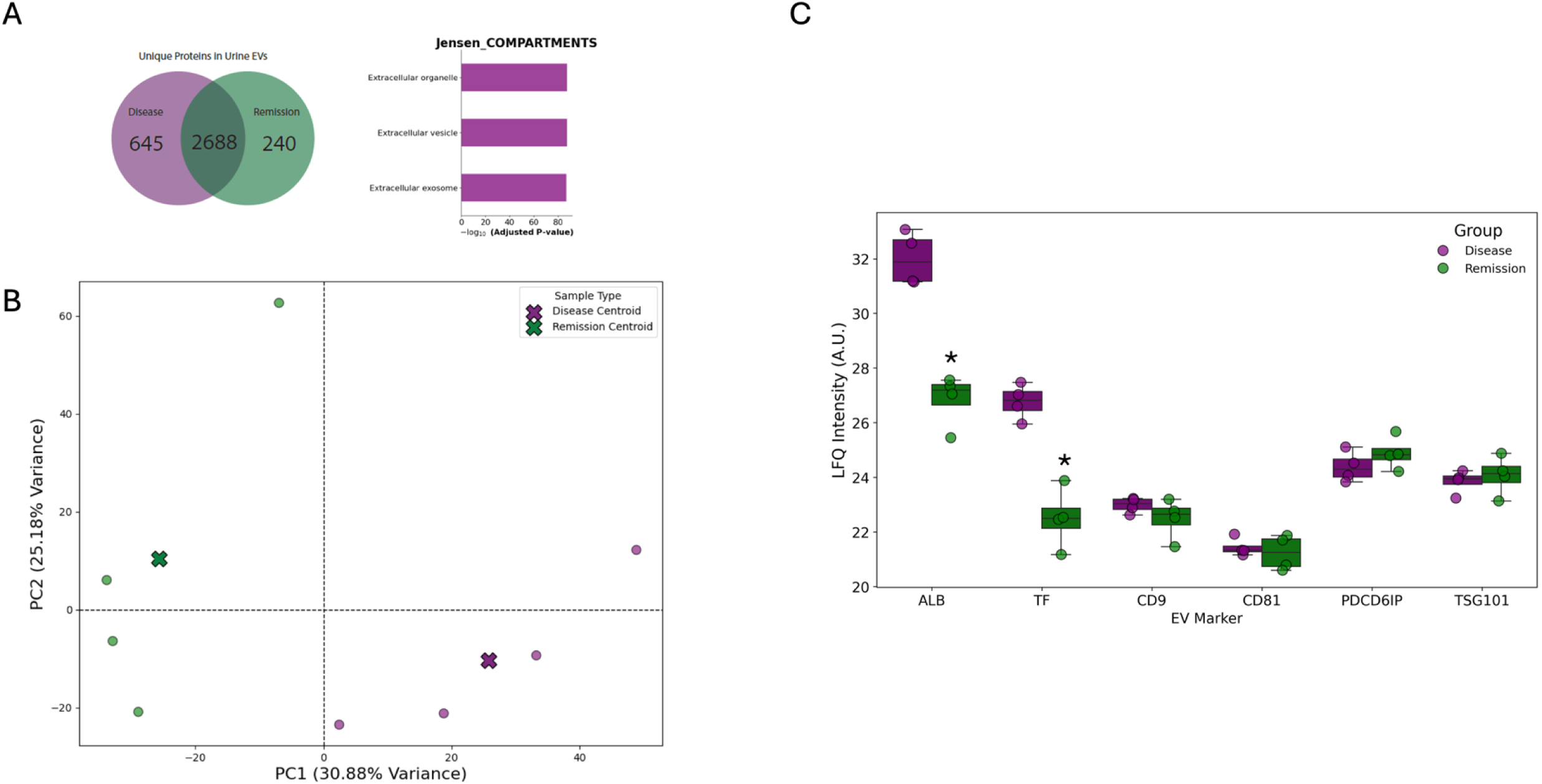
A. Venn diagram showing overlapping and distinct differentially expressed proteins (DEPs) across human nephrotic syndrome (NS) states. Disease, referring to active NS and proteinuria, versus remission, where there is no longer active disease. These are DEPs specific to urinary large extracellular vesicles (LEVs) taken from clinical specimens. We identified 240 DEPs specific to remission, 645 specific to disease, and 2688 overlapping DEPs. Jensen compartment analysis reveals that the LEV DEPs identified are largely specific to extracellular organelle, vesicle, and exosome. B. Principal component analysis comparing LEVs from urinary specimens between disease and remission conditions. C. Bar chart showing comparisons of specific EV proteins across disease and remission conditions. There was enrichment in albumin and tissue factor in disease EVs compared to remission. This likely reflects disease status. There were no differences between disease and remission in relation to common EV markers, including CD9, CD81, Programmed Cell Death 6 Interacting Protein (PDCD6IP; also known as ALIX), and Tumour Susceptibility Gene 101 (TSG101). This suggests a uniform isolation of EVs across each disease condition. *, P<0.05.

An in-depth analysis of urinary LEV proteomes identified 512 DEPs being upregulated in patients with active NS disease compared to 350 DEPs downregulated in patients in remission (**Figure 5A**). GSEA identified significant enrichment of Solute Carrier Membrane (SLC) transport proteins under disease conditions (**Figure 5B, C**). Given that these proteins are expressed throughout the nephron, their enrichment likely reflects disease status. In contrast, remission LEVs were enriched in epidermal growth factor (EGF) and COL6A1, two proteins with known functions in tissue regeneration [14]. Collectively, these findings support our hypothesis that the proteomes of LEVs reflect disease status and that several enriched proteins likely have tissue-specific origins.

**Figure 5.**
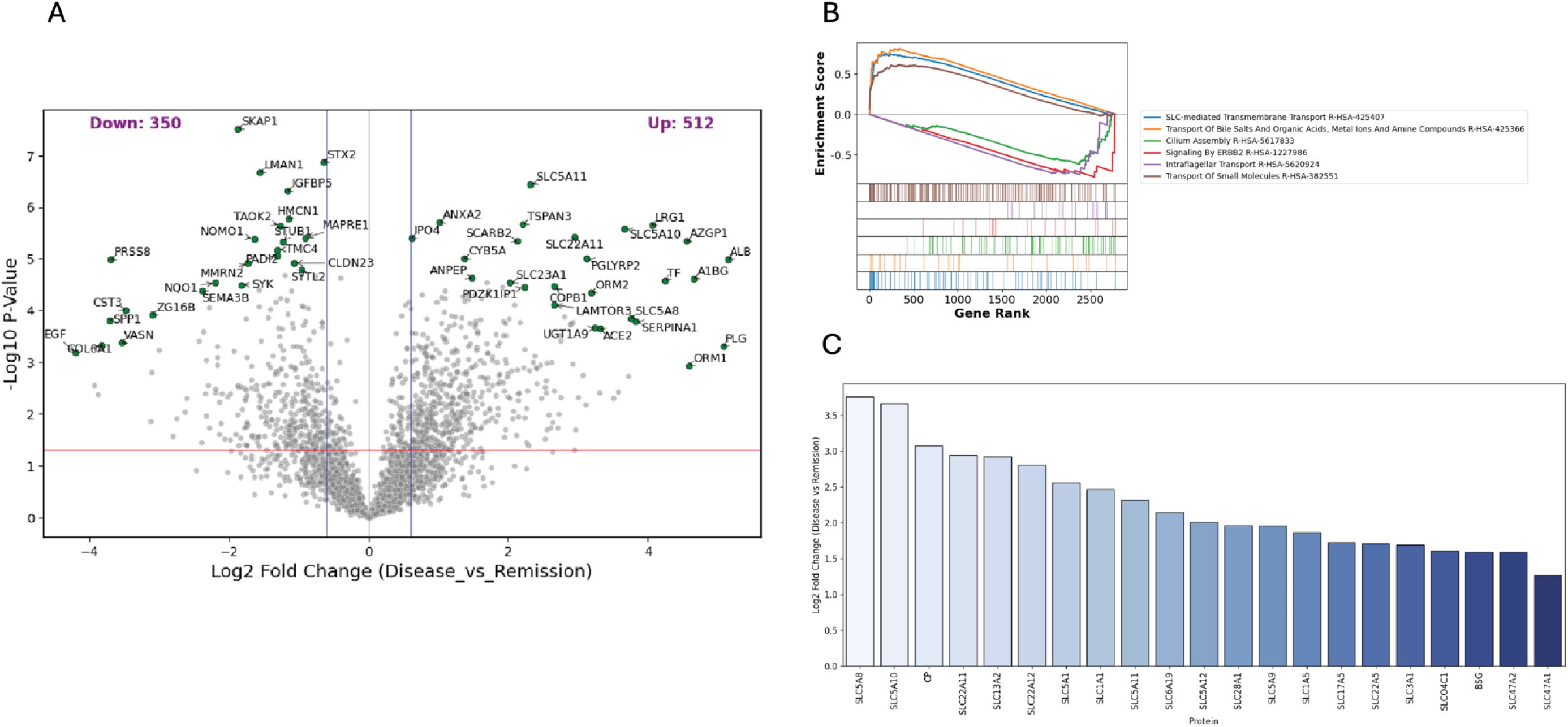
A. Volcano plot comparing large extracellular vesicle (LEV) differentially expressed proteins (DEPs) from urinary specimens from children with nephrotic syndrome (NS) between disease and remission states. We identified 512 upregulated DEPs in disease LEVs compared with 350 downregulated DEPs, relative to remission. B. Gene set enrichment analysis revealed upregulated DEPs including Solute Carrier (SLC) transporters, intraflagellar transport, and transport of small molecules. C. Bar chart showing multiple upregulated DEPs from the SLC transporter family.

### There are 36 LEV DEPs identified following treatment with PAN that were observed in LEVs from children with NS disease

Following a less biased approach to evaluating urinary LEV DEPs, we sought to align those LEV DEPs identified in hPods following PAN injury (versus control), and DEPs present on LEVs from the urine in children with NS disease (versus remission). Specifically, we wanted to directly compare LEV DEPs between experimental NS in the lab, and clinical NS LEVs during disease. We identified 36 significantly altered DEPs under both conditions. There were commonalities between experimental NS and human NS, specifically Syntaxin 3 (STX3), Regulatory Particle Non-ATPase 2 (RPN2), and Dynactin Subunit 1 (DCTN1) were all commonly upregulated (**Figure 6**). Conversely, Heat Shock Protein Family A member 9 (HSPA9), and phosphate cytidylyltransferase 2, ethanolamine (PCYT2) were downregulated. We did however note certain differences in protein abundance between experimental and human NS. In this regard, Prothrombin (F2), and Complement C7 (C7) were opposite, with F2 being upregulated in NS disease LEVs, but downregulated in experimental NS, similar to C7. LEV C9 was markedly upregulated under NS disease conditions, but in experimental NS, it was mildly downregulated.

**Figure 6.**
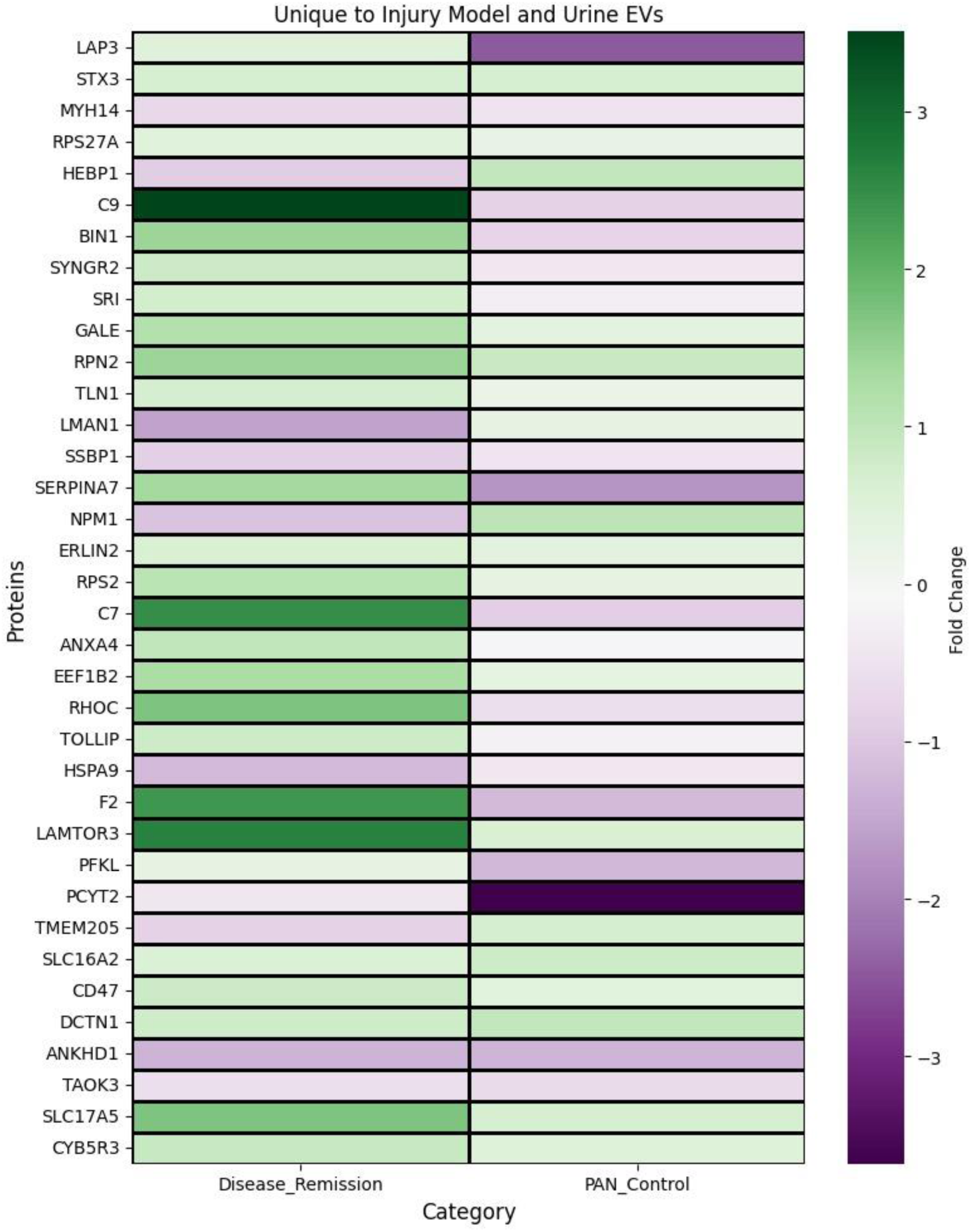
A. Heatmap comparing fold-change in differentially expressed proteins (DEPs) from large extracellular vesicles (LEVs) taken from experimental NS (puromycin aminonucleoside (PAN) versus control) and compared to clinical samples (disease versus remission). Fold change is highlighted in green for increases, and purple for decreases. This bar chart allows for direct comparison of fold change increases or decreases in proteins of interest between experimental NS (PAN versus control) and clinical NS (disease versus remission).

## Discussion

The overarching goal of this study was to characterize proteomic changes in a cell culture ‘experimental’ model of NS (the PAN model) in comparison to urine from children with NS. In our *in vitro* model, we observed that following injury with PAN, there were many DEPs in hPods, as well as in their LEVs. We observed DEPs in urinary LEVs from children with NS disease versus remission; however, many of these DEPs were not specific to the podocytes yet were reflective of the global microenvironment of the injured nephron. Notably, we were able to identify 36 DEPs which were shared between LEVs from PAN-injured podocytes and LEVs from patients with NS disease. To our knowledge, this is the first study to describe proteomic DEPs in LEVs between children with NS disease relapse and remission and relate these DEPs to an *in vitro* podocyte model.

We identified TP53 as a protein of interest in hPods treated with PAN. The role of TP53 is multifaceted, as it is involved in cell death pathways, and can also act as a master cell cycle regulator, and is involved in DNA repair, and senescence [15]. In the presence of PAN, we speculate that TP53 is upregulated to counteract cellular stressors, such as oxidative stress and DNA damage. Additionally, Cyclin-Dependent Kinase Inhibitor 1A (CDKN1A; p21) was upregulated in PAN-injury hPod LEVs, likely reflecting cellular stress [15]. We also observed upregulated proteins associated with cilia and flagella (Cilia and Flagella Associated Protein 20; CFAP20) and actin filaments (Tropomodulin 2; TMOD2), which is not surprising given that the podocyte response to cellular stress includes effacement [16]. To our knowledge, this is the first observation of such proteins being altered in an hPod model.

Moving beyond the limitation of the hPod model, we examined urinary LEVs from children with NS. A novel observation from this study includes the discovery that several tubular proteins are upregulated in urinary LEVs during times of disease relapse in children with NS. We observed significant enrichment of SLC Transporters, which localize mostly to the kidney tubules, and have been described in urinary EVs in the past. They are also present in podocytes, as outlined by Prunotto *et al*., where they analyzed the proteome of podocyte EVs from normal human urine and identified SLC-12A1, −12A3, −13A2, and −23A1 [17]. Consistent with this, we identified SLC13A2 as being upregulated in disease-relapse urinary LEVs. When comparing our results to a study by Gonzales *et al.,* which evaluated total urinary EVs, we see similar trends with a more marked increase in SLC Transporters (including SLC5A8, SLC5A10, and SLC22A5) [18].

Lastly, we sought to identify podocyte-specific LEV proteomic signatures by comparing our *in vitro* hPod data (PAN versus control) with NS disease LEV data (disease versus remission). In doing so, we identified 36 overlapping, differentially regulated DEPs. Importantly, there were commonly up- and down-regulated proteins across both experimental conditions. STX3 and DCTN1 were identified as upregulated DEPs in both experimental NS as well as in NS disease LEVs. STX7, another of the Syntaxin family, has been found abundantly expressed in podocyte clathrin-coated vesicles [19]. Similarly, DCTN1 was also identified in this same study. We speculate these proteins, among many of the other upregulated DEPs in LEVs represent responses to stress or vesiculation.

Our paper is not without its limitations. As discussed extensively, the hPod model is simply a model and cannot definitively recapitulate the glomerular environment [20]. As such, we must interpret these data with caution and use them to guide future studies. A viable next step might include interrogation of directly differentiated podocytes from pluripotent stem cells, or rodents, treated with anti-podocyte antibodies [21]. Another limitation is that the patient-derived urinary LEVs were not podocyte-specific but were a mix of LEVs originating from multiple cellular sources, including tubular. This underscores the need for examination of the tubular cell LEV proteome *in vitro*, which is a future direction for our lab.

Strengths of this current paper include the ‘ground up’ approach taken to interrogate the impact of PAN on hPods *in vitro*, in the context of its use as an experimental NS model. Next, through evaluation of pure hPod LEVs, and comparing them to urinary-derived LEVs from children with NS, we attempted to determine truly podocyte specific LEVs in a mixed urinary milieu. The main limitation with this analysis is the fact that proteins discovered in the hPod LEV proteome may be present in non-podocyte kidney cells, like tubular cells. With this limitation in mind, we did observe differential abundance of proteins across *in vitro* conditions and the human LEVs. Several of the identified proteins have been reported in podocytes, both *in vitro* and *in vivo*, thus suggesting that perhaps some of our upregulated NS LEV DEPs may be specific to podocytes. Further validation of this will be paramount; and determining their importance in childhood NS is the obvious next step.

In summary, hPod LEVs are increased *in vitro* following PAN injury. This supports and builds upon our findings of increased podocyte urinary LEVs from children with NS. On further interrogation, podocytes *in vitro* have distinct proteomes when exposed to PAN. Several of the identified DEPs in our study have been described previously in podocytes and potentially have important functions therein. By isolating LEVs from the urine of children with NS during both disease and remission, we have developed the first urinary LEV proteome specific to this condition in children. Uncovering the role of these up- and down-regulated DEPs in podocytes and elucidating the significance of their inclusion in LEVs are clear next steps.

The authors declare no conflicts of interest exist.

---

**Supplemental Figure 1.**
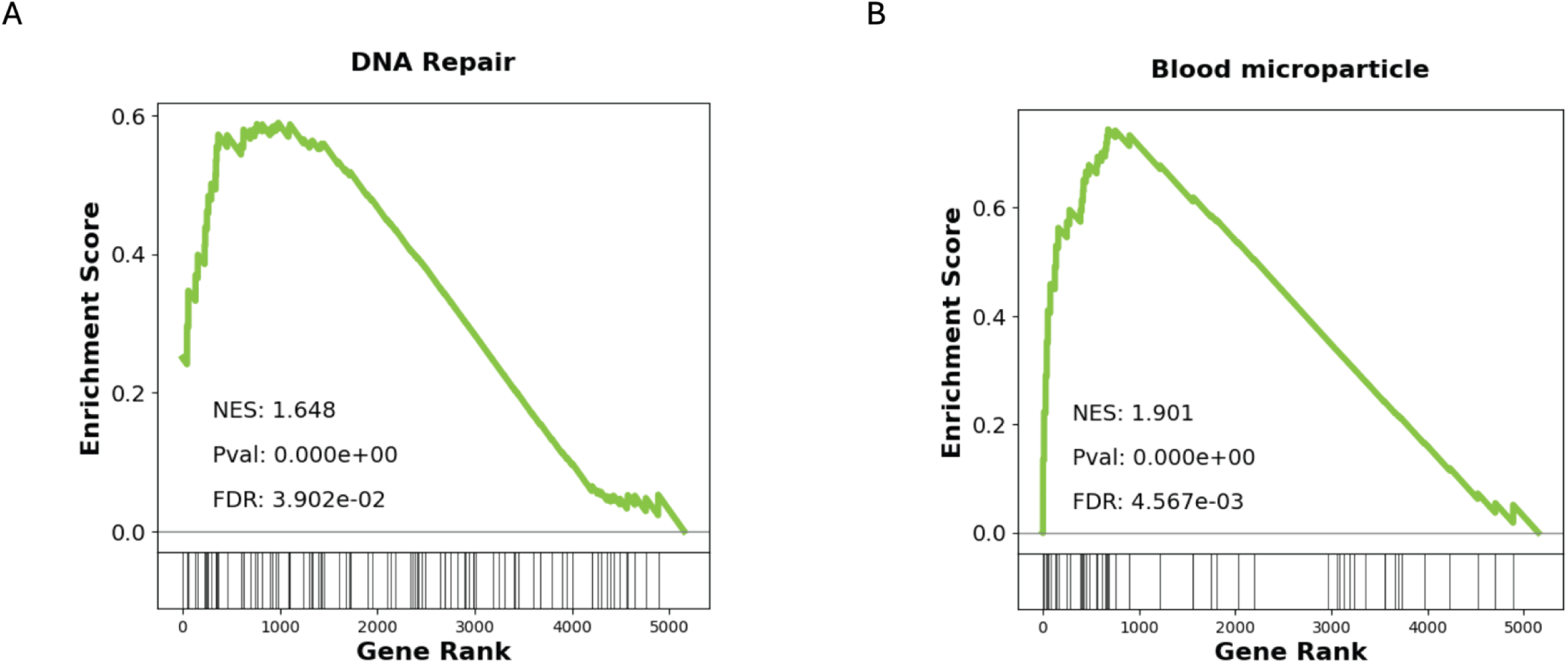
A. Gene set enrichment analysis (GSEA) from cellular proteome analysis, comparing puromycin aminonucleoside (PAN) treatment with control. GSEA analysis revealed that PAN treatment, relative to control, led to an increase in proteins associated with DNA repair. B. Similar to A, GSEA data from PAN-treated versus control hPods revealing that PAN led to an increase in proteins associated with microparticles (previous nomenclature for large extracellular vesicles, LEVs).

**Supplemental Figure 2.**
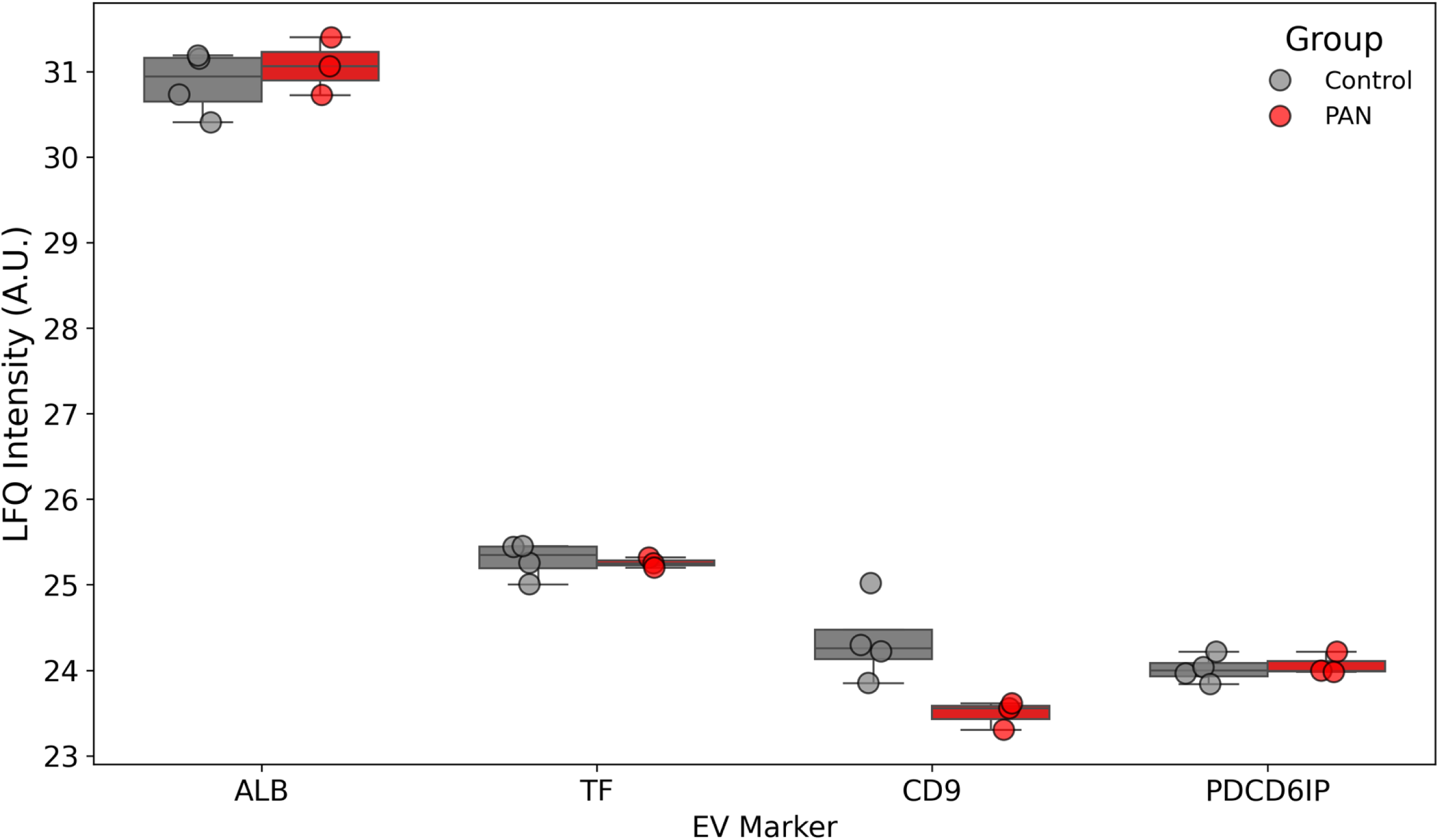
A. Bar chart showing comparisons of specific EV proteins across each experimental condition. No significant differences were observed across each experimental condition for EV-related albumin, tissue factor, CD9 and, Programmed Cell Death 6 Interacting Protein (PDCD6IP; also known as ALIX). This suggests a uniform isolation of EVs across each experimental condition.

**Supplemental Figure 3.**
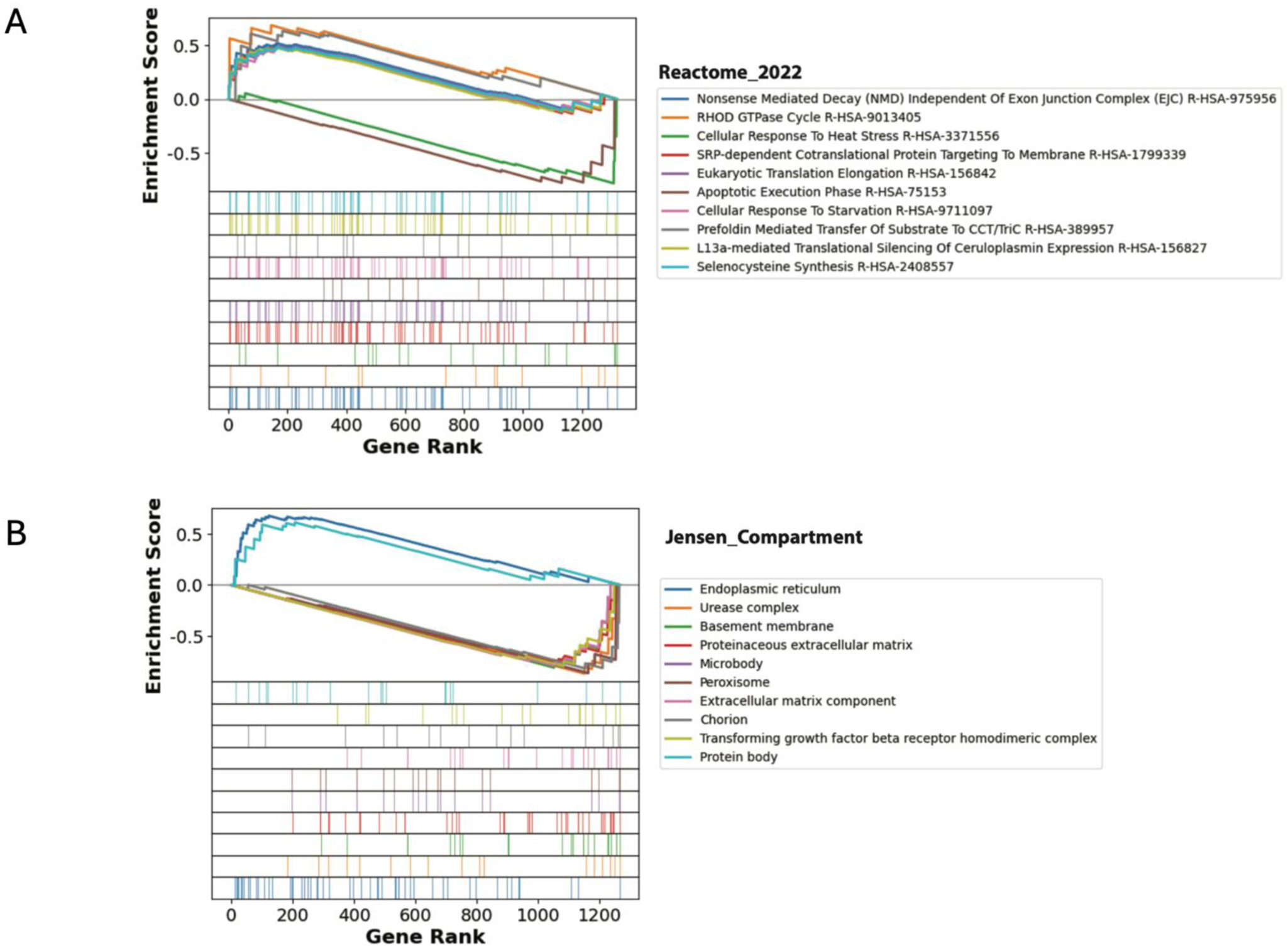
A. Gene set enrichment analysis (GSEA; Reactome 2022) for hPod large extracellular vesicles (LEV) comparing PAN versus control experimental conditions. B. Similar to A; however, using Jensen Compartments.

**Supplemental Figure 4.**
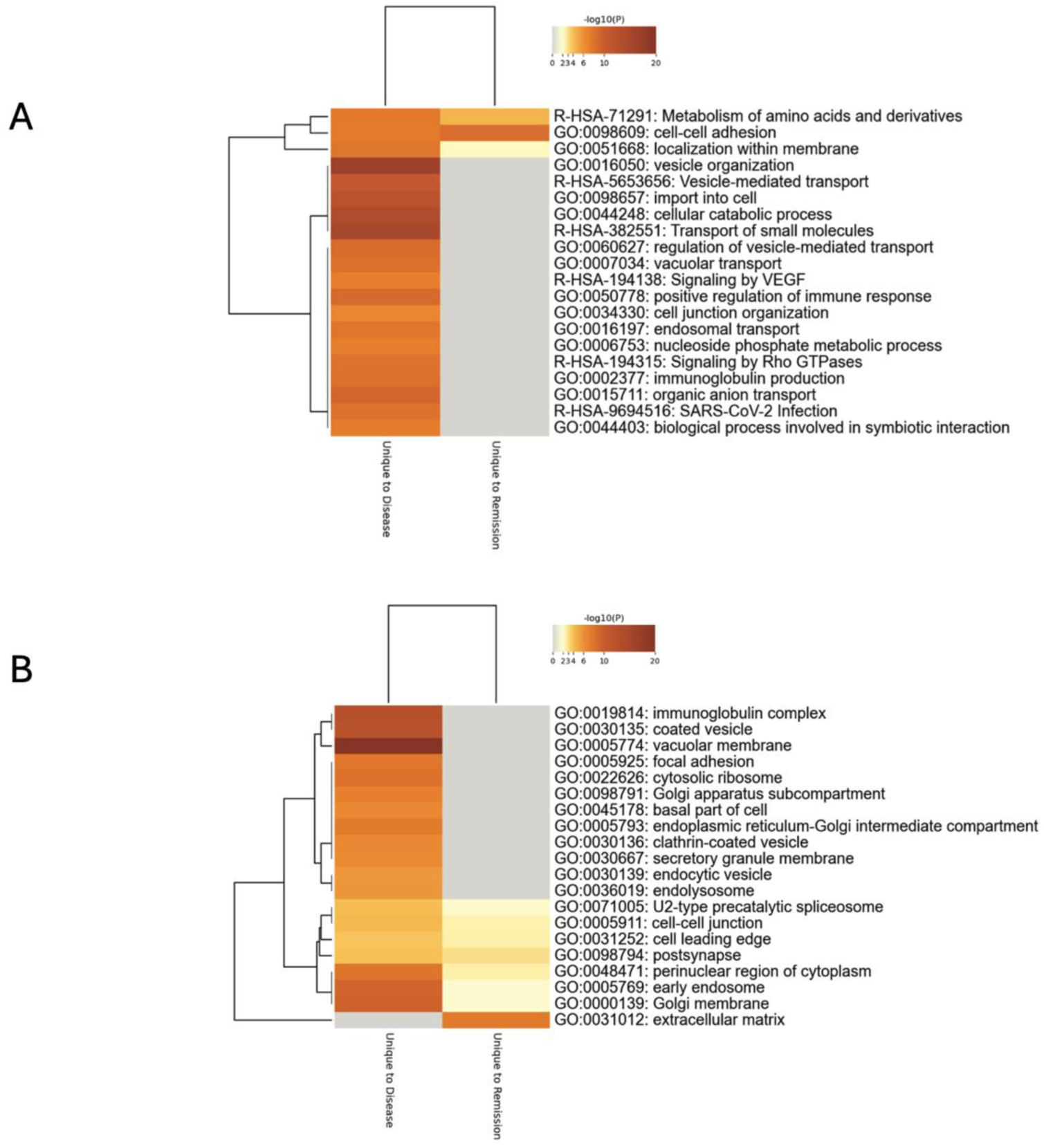
A. Heat map (Reactome, Gene Ontology (GO)) comparing urinary large extracellular vesicle (LEV) differentially expressed proteins (DEPs) between clinical specimens taken from children with nephrotic syndrome (NS), comparing those with disease versus remission. B. Similar to A; however, using only GO terminology.

**Supplemental Table 1.**

|  | **Relapse or remission** | **Steroid profile** | **Sex<sup>#</sup>** | **Age** | **Urine protein:creatinine ratio (g/mmol)** | **Biopsy?** |
| --- | --- | --- | --- | --- | --- | --- |
| **Patient 1** | Relapse | Sensitive | M | 15 | 2.29 | *^ |
| **Patient 2** | Relapse | Resistant | M | 10 | 0.38 | ** |
| **Patient 3** | Relapse | Sensitive | M | 3 | 2.14 | *^ |
| **Patient 4** | Relapse | Resistant | F | 8 | 0.63 | *^ |
| **Patient 5** | Remission | Resistant | M | 16 | 0.01 | *** |
| **Patient 6** | Remission | Sensitive | M | 5 | 0.02 | *** |
| **Patient 7** | Remission | Resistant | M | 4 | 0.02 | *** |
| **Patient 8** | Remission | Sensitive | M | 9 | 0.01 | *** |

\* No biopsy.
\*\* Focal segmental glomerulosclerosis lesions.
\*\*\* Minimal change lesions.
^ Presumed minimal change lesions (classic idiopathic NS in children is presumed to associate with minimal change lesions on biopsy, and thus in steroid sensitive children, a biopsy is not always performed).

